# Pharmacological recruitment of a VTA glutamatergic arousal circuit by the natural product BN3 drives broad-spectrum emergence from general anesthesia

**DOI:** 10.64898/2026.02.17.706054

**Authors:** Ran Ji, Yi Zhao, Zhenbo Huang, Xuehan Li, Yuhao Chen, Yu Gan, Tian Zhang, Ao Hai, Shirong Lai, Tianyou Liu, Yudong Yin, Yuhao Sun, Yixin Yuan, Buyi Xu, Jin Liu, Bowen Ke

## Abstract

Pharmacological control of emergence from general anesthesia remains strikingly limited. Clinically available antagonists are largely drug-class specific, leaving recovery dependent on anesthetic clearance rather than to an actively programmable switch in brain state. Through phenotypic screening of a structurally diverse natural-product library, we identified BN3 (L-(−)-camphor) as a broad-spectrum emergence agent. BN3 robustly shortened recovery from propofol anesthesia across species—from mice and rats to rabbits, dogs, and rhesus monkeys—and remained effective after continuous infusion. In rodents, BN3 also accelerated emergence from mechanistically distinct anesthetics, including etomidate, remimazolam, sodium γ - hydroxybutyrate, midazolam, ketamine, sodium pentobarbital, and isoflurane. EEG recordings revealed a rapid cortical shift from slow-wave dominance to a wake-like high-frequency spectrum. Mechanistically, BN3 preferentially recruited ventral tegmental area (VTA) glutamatergic neurons: chemogenetic silencing abolished the BN3-evoked EEG spectral shift and strongly attenuated behavioral acceleration of emergence, whereas optogenetic activation of these neurons was sufficient to promote recovery. BN3 increased glutamate release in the nucleus accumbens and medial prefrontal cortex, and projection-specific inhibition of VTA→NAc or VTA→mPFC outputs blunted BN3-induced emergence. These findings define a VTA-centered glutamatergic arousal circuit that can be pharmacologically recruited as a common gateway for anesthetic-independent emergence.

## Introduction

General anesthesia is routinely described as “reversible,” yet the field has surprisingly little mechanistic control over the moment of reversal [1]. In practice, emergence is typically managed as a pharmacokinetic endpoint: clinicians wait for anesthetic concentrations to fall, while mitigating physiological instability and complications that may accompany delayed or dysregulated recovery [2, 3]. This reliance on clearance reflects a deeper constraint. Although the molecular pharmacology of anesthetic-induced unconsciousness is diverse, the pharmacology of *awakening* is narrow [4]. As a result, emergence remains the least programmable phase of perioperative brain-state control.

This gap is not simply technical; it is conceptual. If anesthetic unconsciousness were reversed only by negating each anesthetic’s molecular actions, then a universal emergence agent would be unlikely—given that commonly used anesthetics act through distinct receptors and network effects [5]. Consistent with this, clinically available reversal strategies are largely class-restricted: a benzodiazepine antagonist can reverse benzodiazepine-based hypnosis, but does not address unconsciousness induced by non-benzodiazepine hypnotics [6], and no broadly effective “off switch” exists for general anesthesia across regimens [7]. The absence of such an agent has reinforced the notion that emergence is fundamentally passive.

However, a growing circuit-level view argues otherwise: awakening is an active brain-state transition that can be driven by endogenous arousal systems, and may be mechanistically dissociable from induction [8]. In this framework, anesthetics may converge on a limited set of network configurations that stabilize unconsciousness, whereas emergence reflects the re-engagement of conserved arousal pathways capable of destabilizing that configuration. If so, the most general route to accelerating emergence may not be receptor-level antagonism of each anesthetic, but rather pharmacological recruitment of a shared arousal circuit that functions as a common gateway back to wakefulness. The key challenge is to identify small molecules that can engage such circuitry with sufficient potency and generality to override anesthetic diversity—and to map the circuit mechanism with causal resolution.

Natural products offer privileged chemical space for this search [9]. Their structural diversity and historical enrichment for neuroactive scaffolds make them well suited for phenotypic approaches that prioritize state transitions over presumed molecular targets [10, 11]. We therefore constructed a structurally diverse natural-product compound library and performed a phenotypic screen for agents that shorten time to emergence following a standardized propofol challenge, identifying BN3 (L-(−)-camphor) as a leading hit.

Here we show that BN3 functions as a broad-spectrum emergence agent across anesthetics and species. BN3 accelerates behavioral recovery from various anesthetics with distinct mechanisms— including propofol, etomidate, remimazolam, sodium γ-hydroxybutyrate, midazolam, ketamine, sodium pentobarbital, and isoflurane in rodents. More importantly, its efficacy extends from rodents to rabbits, dogs and rhesus monkeys and persists under continuous-infusion paradigms. BN3-driven behavioral recovery is accompanied by a rapid EEG shift toward a wake-like spectral profile, indicating a genuine reconfiguration of brain state rather than a purely motor effect. Mechanistically, we identify a ventral tegmental area (VTA)-centered glutamatergic circuit as both necessary and sufficient for BN3-induced emergence: BN3 rapidly activates VTA glutamatergic neurons, inhibition of this population abolishes BN3’s effects, and projection-specific suppression of VTA glutamatergic outputs to the nucleus accumbens or medial prefrontal cortex prevents BN3 from accelerating recovery. Together, these data establish a circuit-based strategy for anesthetic-independent reanimation and nominate BN3 as a tractable chemical entry point to the neural control of emergence.

## Methods

### Mice

All experimental protocols were approved by the Institutional Animal Care and Use Committee of Sichuan University West China Hospital (protocol no.:20251011001). *C57BL/6J* (000013) mice were obtained from GemPharmatech and *Vglut2-ires-Cre: Slc17a6tm2(cre)Lowl/MwarJ* (028863) mice were obtained from The Jackson Laboratory. All mice used in the experiments were male and aged 8weeks at the start of the stereotaxic injections and experiments. Mice were maintained on a 12-h light/12-h dark cycle in a controlled environment with constant temperature and humidity, and with free access to food and water.

### Drugs

BN3 was dissolved in a medium- and long-chain triglyceride emulsion (C6–C24) at a concentration of 12 mg/ml and sonicated at 37°C for 15 minutes. The subsequent experimental dose was 48 mg/kg. Propofol (22.11 mg/kg), etomidate (3.36 mg/kg), remimazolam (57.35 mg/kg), sodium γ-hydroxybutyrate (1300 mg/kg), flumazenil (3 mg/kg), midazolam (50 mg/kg) and ketamine (80 mg/kg) were all administered intravenously. Sodium pentobarbital (94.5 mg/kg) was administered intraperitoneally. The effect of BN3 on inhaled anesthetics was tested after mice were anesthetized for 30 minutes with 1.5% isoflurane (RWD Life Technology Co., Ltd, Shenzhen, China).

### Virus preparation

All viruses were purchased from BrainVTA technology Co., Ltd. (Wuhan, China) or BrainCase Co., 169 Ltd. (Shenzhen, China), including rAAV-hSyn-hM4D(Gi)-EGFP-WPRE-hGH pA (5.04×10^12^ genome copies per ml, 200nl, BrainVTA), rAAV-hSyn-GCaMp6s-WPRE-hGH pA (5.22×10^12^ genome copies per ml, 200nl, BrainVTA), rAAV-hSyn-DIO-ArchT-EGFP-WPRE-hGH pA (5.04×10^12^genome copies per ml, 200nl, BrainVTA), rAAV-hSyn-DIO-GCaMp6s-WPRE-hGH pA (5.06×10^12^genome copies per ml, 200nl, BrainVTA), rAAV-EF1α-DIO-hM4D(Gi)-EYFP-WPRE-hGH pA (4.50×10^12^genome copies per ml, 200nl, BrainVTA), rAAV-EF1α-DIO-hChR2(H134R)-EYFP-WPRE-hGH pA (5.08×10^12^genome copies per ml, 200nl, BrainCase), AAV-hsyn-iGluSnFR3.v857-WPRE-PA (5.32×10^12^genome copies per ml, 200nl, BrainVTA).

### Stereotaxic injections

Throughout the entire surgical procedure, mice were anesthetized with 2% isoflurane and secured in a stereotaxic frame (RWD Life Technology Co., Ltd, Shenzhen, China), with body temperature maintained at a stable level throughout the operation. To prevent dry eyes, eye ointment was applied preoperatively. Hair was shaved and the area sterilized with povidone-iodine solution. The skin was incised to expose the skull, and the periosteum was removed. A skull drill was then used to create a 0.5 mm diameter hole above the target injection site. Finally, 200 nl of virus was injected into the target area using a micropipette (Hamilton, catalog no. 7002KH).

Stereotaxic coordinates for virus injection and optic fiber implantation (bregma/lateral/dorsoventral in mm):

VTA: AP -3.3 mm, ML 0.13 mm, DV -3.93 mm.

PVT: AP -0.3 mm, ML 0.00 mm, DV -3.3 mm.

NAc: AP 1.1 mm, ML 0.60 mm, DV -4.2 mm.

mPFC: AP +1.78 mm, ML ±0.25 mm, DV -2.5 mm.

### Immunohistochemistry and fluorescence in situ hybridization (FISH)

After deep anesthesia, the mice were perfused with PBS, followed by a 4% paraformaldehyde solution. Brains were extracted and post-fixed in 4% paraformaldehyde for 24 hours, then placed in 30% sucrose PBS at 4°C until tissues settled to the bottom. For cryopreservation, the tissue was embedded in an embedding medium. Then, we used a cryostat (Leica) to cut the brain into 15 μm (for FISH) or 30 μm (for immunohistochemistry) coronal sections. For immunohistochemical staining, the sections were washed three times with PBS, incubated in a blocking solution (1% bovine serum albumin) for two hours and then incubated overnight at 4°C with a primary antibody (rabbit anti-c-Fos, 2250S, 1:800, Cell Signalling Technology). The following day, the sections were washed three times in TBS buffer (0.4%Triton X-100 in PBS) and incubated with the secondary antibody (goat anti-rabbit Alexa-488, A21206, 1:500, Thermo Fisher Scientific) for two hours. FISH experiments were performed using the RNAscope kit according to the manufacturer’s instructions (Advanced Cell Diagnostics). Fluorescent images were acquired using a Zeiss LSM980 confocal microscope and a Nikon Eclipse 80i fluorescence microscope.

### Electroencephalogram Recording and Analysis

Mice were placed in a 2% isoflurane induction chamber and anesthetized before being transferred and secured in a stereotaxic apparatus. Electrodes were fixed to the skull surface and sealed with Denture Base Materials. Mice were allowed to recover for at least 7 days post-surgery. On the recording day, mice were first connected to the EEG system and acclimated for 20 minutes in the recording cage at 25°C, allowing free movement. Following adaptation, baseline EEG was recorded for 2 minutes. After intravenous administration of anesthetic, recording continued for 1 minute. Following intravenous administration of BN3, recording proceeded until the mouse recovered. All EEG signals were recorded using Pinnacle’s three-channel tethered EEG/EMG systems (Pinnacle Technology, USA) at a sampling rate of 400 Hz. Signals were filtered within the 0.5–30 Hz frequency range.

### Fiber Photometry Recording

To detect neuronal calcium signals, rAAV-hSyn-GCaMp6s-WPRE-hGH pA or rAAV-hSyn-DIO-GCaMp6s-WPRE-hGH pA was injected to VTA. To assess glutamate dynamics, AAV-hSyn-iGluSnFR3.V857-Wpre-PA was injected into target brain regions (NAc and mPFC). Surgical details are described below: the animals were injected with the virus while under anesthesia with 1.5% isoflurane, two weeks after the viral injection, a fiber (core diameter: 200 µm, ThinkerTech, China) was implanted above the injection site and secured with dental acrylic resin. The mice were given a week to recover. Three weeks after viral injection, fluorescent signals were collected using the fiber recording system. Prior to the recording days, the mice underwent a 20-minute adaptation period in the recording cage. The fiber was connected to an optical fiber photometry system (ThinkerTech, China). The excitation of the calcium sensor or glutamate sensor was achieved using a 470 nm LED, with a 405 nm LED serving as an isochromatic reference light source. The laser parameters were as follows: wavelength 470 nm and power 10–20 mW. Signals were downsampled to 40 Hz for analysis. All recordings were conducted from 1 min before propofol administration to 1 min after BN3 administration, with the time point of BN3 administration defined as 0. The analyzed data included fluorescence values from one minute after propofol administration (from -1 to 0 minutes) and one minute after BN3 administration (from 0 to 1 minutes). The area under the curve (AUC) values were restricted to time window 0-s to 50-s after BN3 delivery for GCaMp6s signals and to time window 0-s to 30-s after BN3 delivery for iGluSnFR3 signals. The percentage Δ*F/F* was calculated by 100 × (*F* − *F*_mean_)/*F*_mean_, where *F*_mean_ was the mean fluorescence intensity. The Z-score was calculated using MATLAB by

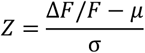

Data analysis was performed using custom MATLAB scripts (MATLAB R2021a).

### Optogenetic manipulation

The fibers were implanted into the ventral tegmental area (VTA) two weeks following the viral injection. The surgery was performed under 2% isoflurane anesthesia and involved drilling a hole and inserting an optical fiber (core diameter: 200 µm, ThinkerTech, China) into the target area. Once the target depth was reached, the fiber was secured with dental cement. The animals then underwent a one-week recovery period prior to the start of experiments. Optogenetic activation experiments employed blue light stimulation (473 nm, 2–5 mW, 15 ms pulses at 20 Hz). For optogenetic silencing experiments, continuous green light stimulation (wavelength 580 nm, power 5–8 mW) was employed. All control groups underwent the same experimental procedures and light stimulation.

### Chemogenetic manipulation

To achieve chemogenetic modulation of neuronal activity, we expressed CNO-based inhibitory DREADDs (hM4Di) via an adeno-associated virus vector. On day 21 post-AAV microinjection, CNO (MedChemExpress, HY-17366, 3 mg/kg) was administered intraperitoneally 30 minutes before propofol and BN3 to activate the DREADDs, with an equal volume of saline serving as the control.

### Quantification and statistics

All statistical analyses were performed using GraphPad Prism 9.0 (GraphPad Software, Inc., USA). For comparisons between two groups, paired or unpaired two-tailed Student’s *t*-tests were used as appropriate. For comparisons among multiple groups, one-way ANOVA followed by Dunnett’s multiple comparisons test was applied. Detailed statistical results are presented in the figures. Statistical significance was set at *p* less than 0.05. Unless otherwise stated, data are presented as mean ± SEM.

## Results

### 1. BN3 Consistently Shortens Recovery Time from Propofol Anesthesia Across Species

Natural products represent an important resource for drug discovery, with Chinese herbal medicines having produced valuable achievements—most notably artemisinin—within this field. Many Chinese medicinal formulas have been reported to promote arousal in patients with disorders of consciousness in various clinical scenarios [12-14]. To discover novel compounds for promoting arousal, we constructed a structurally diverse natural product compound library and evaluated their potential to accelerate recovery from anesthesia through phenotypic screening (Figure 1A-B, supplementary table 1, supplementary figure 1). Among a series of natural active ingredients, L-(−)-camphor (BN3)—an aromatic organic compound extracted from the leaves and branches of the camphor tree—was identified to possess a remarkable ability to promote recovery from anesthesia (Figure 1B, vehicle: 147 ± 7 s; BN3: 22 ± 4 s; *F*_(12,26)_ =9.09, *p*<0.001). To further assessing the emergence-promoting effect of BN3, we first validated its efficacy in small rodents.

**Figure 1.**
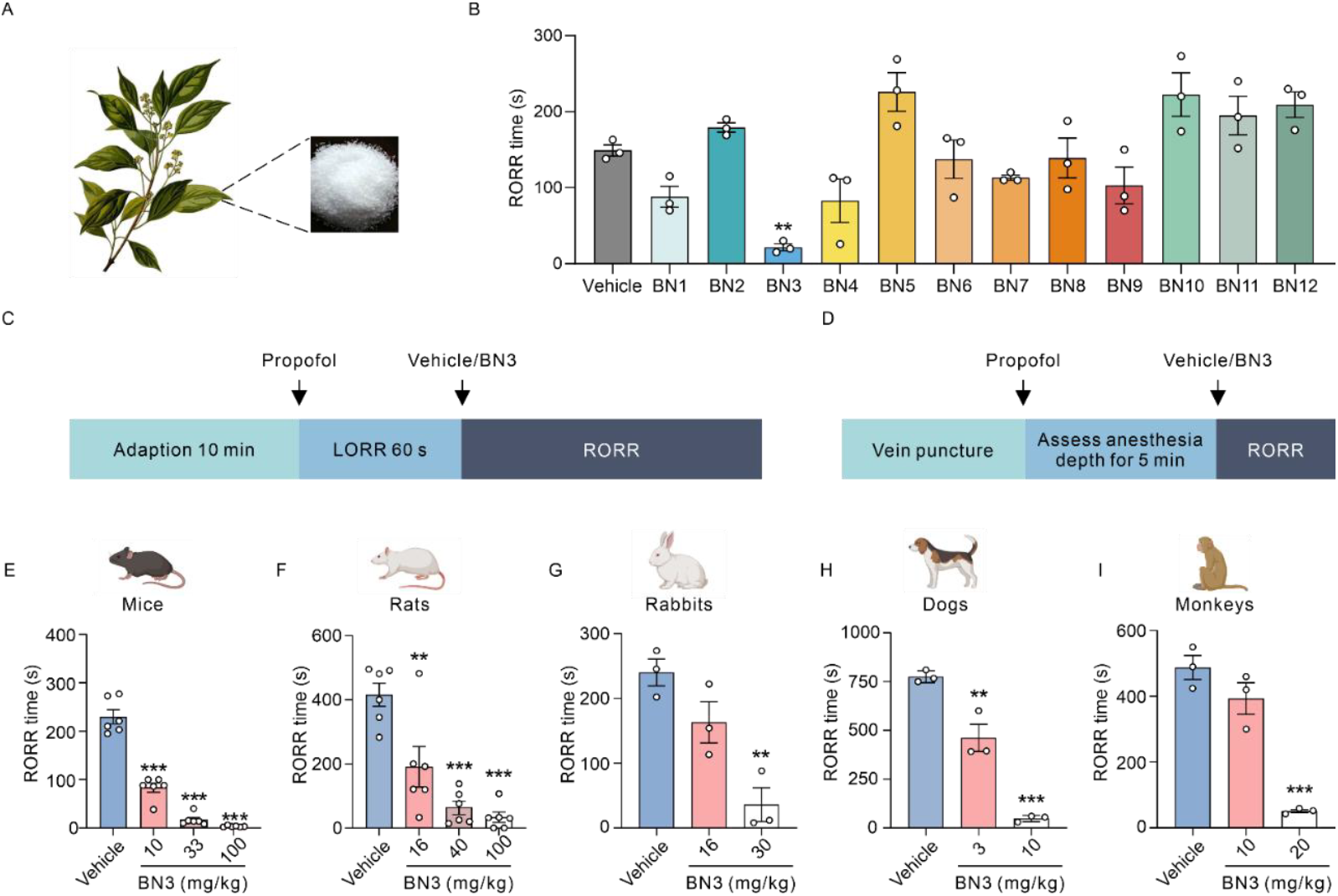
BN3 accelerates emergence from propofol-induced anesthesia across multiple species. (A, B) Screening of natural compounds for their ability to reduce the time to emergence following a standardized propofol bolus in mice. BN3 was identified as the lead compound, n = 3 per group. One-way ANOVA, \*\**p* < 0.01 (C, D) Schematic timelines of the experimental protocols. (C) Protocol for mice, rats, and rabbits. (D) Protocol for dogs and monkeys. LORR: Loss of Righting Reflex; RORR: Return of the righting reflex. (E–I) Dose-response effects of BN3 on time to RORR in (E) mice (n = 6 per group), (F) rats (n = 6 per group), (G) rabbits (n = 3 per group), (H) dogs (n = 3 per group), and (I) monkeys (n = 3 per group). One-way ANOVA, \*\**p* < 0.01, \*\*\**p* < 0.001.

In mice, intravenous administration of BN3 after loss of righting reflex significantly shortened emergence time in a dose-dependent manner (Figure 1C,E, vehicle: 230 ± 15 s; 10 mg/kg: 83 ± 9 s; 33 mg/kg: 17 ± 5 s; 100 mg/kg: 3 ± 1 s; *F*_(3,20)_ =131.20, *p*<0.001). Similar results were observed in rats (Figure 1C,F, vehicle: 416 ± 36 s; 16 mg/kg: 191 ± 63 s; 40 mg/kg: 63 ± 21 s; 100 mg/kg: 34 ± 16 s; *F*_(3,20)_ =20.20, *p*<0.001). We next tested BN3 in larger animals. In rabbits, BN3 (30 mg/kg) significantly reduced recovery time after propofol anesthesia (Figure 1C,G, vehicle: 240 ± 21 s; 16 mg/kg: 163 ± 38 s; 30 mg/kg: 36 ± 26 s; *F*_(2,6)_ =15.06, *p*<0.01). In dogs, a relatively low dose (3 mg/kg) was able to significantly shortened recovery time (Figure 1D,H, vehicle: 776 ± 18 s; 3 mg/kg: 462 ± 69 s; 10 mg/kg: 48 ± 9 s; *F*_(2,6)_ =76.93, *p*<0.001).

To assess translational potential, we tested BN3 in rhesus monkeys. BN3 (20 mg/kg) significantly accelerated recovery from propofol anesthesia (Figure 1D,I, vehicle: 488 ± 36 s; 10 mg/kg: 393 ± 49 s; 20 mg/kg: 51 ± 5 s; *F*_(2,6)_ =42.91, *p*<0.001). Given that propofol is typically administered by continuous infusion in clinical practice, we further examined whether BN3 could reverse propofol anesthesia following continuous infusion. In both rats and rabbits, a single intravenous BN3 injection significantly reduced recovery time following propofol continuous infusion (Supplementary figure 2A-B). Moreover, BN3 was able to induce return of the righting reflex during propofol continuous infusion (Supplementary figure 2C). EEG recordings revealed that BN3 administration shifted brain activity toward a wakeful state, as indicated by decreased delta-band power and increased alpha, beta, and gamma activity (Supplementary figure 2D–F). Post-emergence behavioral assessment showed no signs of locomotion, motor coordination, or spatial recognition memory impairments following BN3 administration (Supplementary figure 3).

Together, these results demonstrate that BN3 consistently shortens recovery time from propofol anesthesia across species, supporting its potential for clinical application.

### 2. BN3 Speeds Recovery from Anesthesia Induced by Various Anesthetic Agents

While a variety of anesthetics with distinct molecular targets are used clinically to induce general anesthesia, no single reversal agent has yet been shown to accelerate recovery across different classes of these drugs. In this study, we investigated whether BN3 could promote emergence from anesthesia induced by agents other than propofol.

Dose–response analysis revealed that BN3 accelerated recovery from propofol anesthesia with an effective dose for 50% response (ED_50_) of 11.95 mg/kg (Figure 2A, *F*_(5,42)_ =108.50, *p*<0.001). To assess its broader utility, we conducted analogous dose-response studies with other routinely used clinical anesthetics. BN3 consistently accelerated emergence from anesthesia induced by etomidate (ED_50_: 23.94 mg/kg; Figure 2B, *F*_(5,42)_ =13.79, *p*<0.001), remimazolam (ED_50_: 16.70 mg/kg; Figure 2C, *F*_(5,42)_ =51.37, *p*<0.001), and sodium γ-hydroxybutyrate (ED_50_: 55.23 mg/kg; Figure 2D, *F*_(5,42)_ =49.85, *p*<0.001).

**Figure 2.**
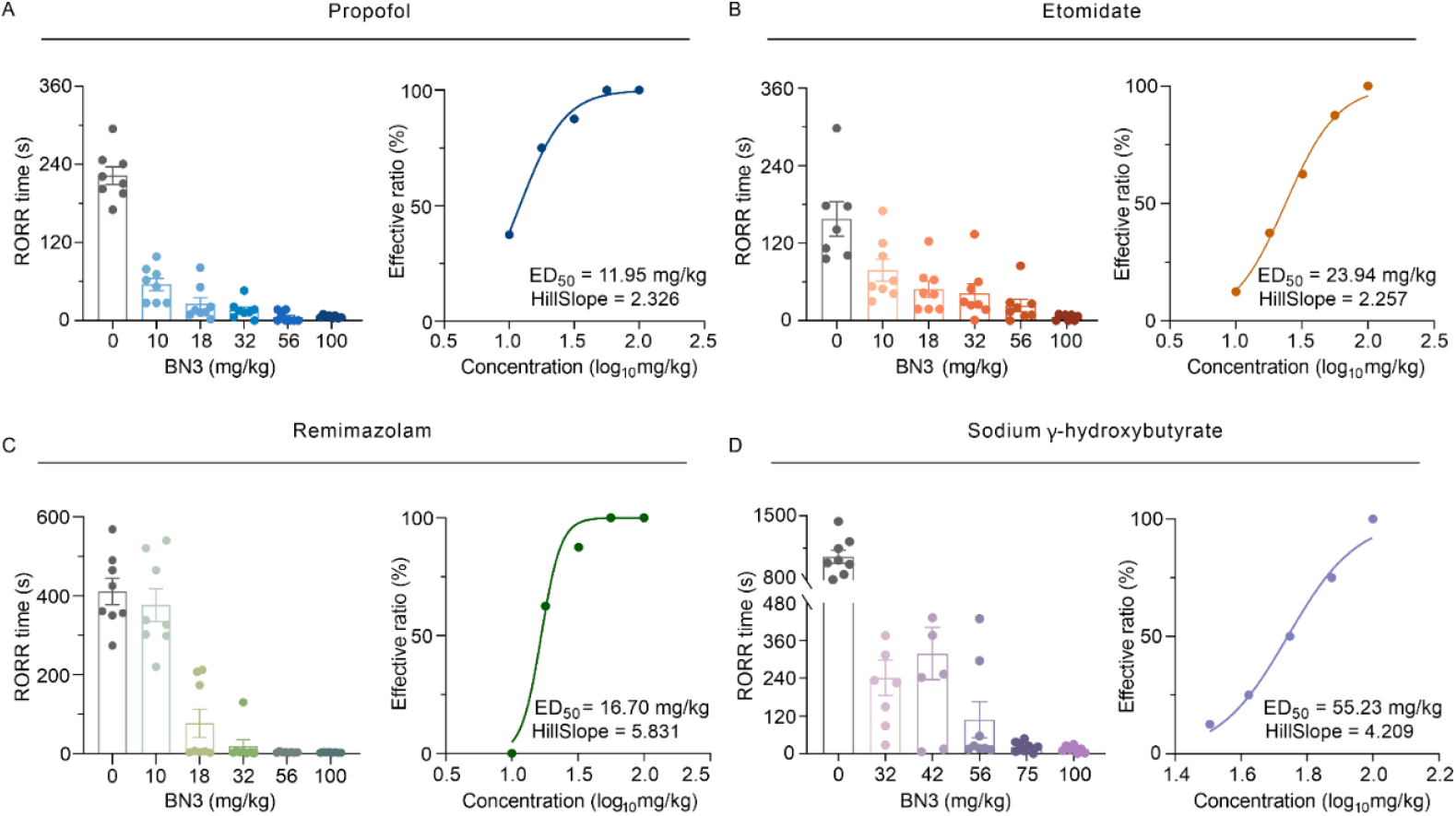
BN3 accelerates emergence from general anesthesia induced by various anesthetics. (A–D) Dose-response curves for the efficacy of BN3 in reducing the time to emergence from (A) propofol, (B) etomidate, (C) remimazolam, and (D) sodium γ-hydroxybutyrate-induced general anesthesia. Mice received BN3 one minute after loss of righting reflex (LORR). The time to return of the righting reflex (RORR) was measured, and data are plotted as percent maximal effect vs. log_10_ dose of BN3. Calculated half-maximal effective doses (ED_50_) are indicated for each anesthetic: (A) 11.95 mg/kg, (B) 23.94 mg/kg, (C) 16.70 mg/kg, and (D) 55.23 mg/kg.

We extended these findings by testing additional anesthetics with diverse molecular targets, including midazolam, ketamine, sodium pentobarbital and isoflurane. In each case, a high dose of BN3 significantly accelerated anesthesia emergence (Supplementary figure 4A-D). For continuous infusion, BN3 also significantly shortened recovery time from etomidate, and remimazolam anesthesia in rats and rabbits (Supplementary figure 4E-F). Finally, we compared BN3 with flumazenil, the clinically approved reversal agent for remimazolam anesthesia. Both BN3 and flumazenil reversed remimazolam-induced anesthesia, but only BN3 effectively reversed propofol-induced anesthesia (Supplementary figure 4G).

Together, these data show that BN3 promotes emergence from anesthesia induced by various anesthetics. Given the structural diversity and distinct molecular targets of these drugs, BN3 is unlikely to act by antagonizing each anesthetic’s molecular mechanism. Instead, BN3 may act via a common, anesthetic-independent pathway. We therefore next explored the neural mechanisms underlying BN3’s emergence-promoting effects, focusing on propofol anesthesia.

### 3. The VTA Is a Key Brain Region Mediating BN3’s Emergence-promoting Effect

Historically, emergence from anesthesia has been viewed as a purely passive phenomenon, occurring gradually as anesthetic drugs are eliminated from the body. Recently, however, it has been suggested that emergence from anesthesia might also accompany an active process that is driven by mechanisms which can be distinct from entry to the anesthetized state. If that is the case, there are two possible ways to reverse the anesthesia process. One way is to target the mechanisms that are responsible for entering the anesthetized state, such as, by antagonizing the molecular targets of anesthesia. The other way is to target the active emergence-promoting process, such as, by activating brain areas promoting wakefulness. Because BN3 promotes awakening from multiple anesthetics with different molecular targets, it is unlikely that BN3 reverses anesthesia by targeting entry mechanisms. Instead, we hypothesized that BN3 activates emergence-promoting brain areas.

To test this, we measured c-Fos expression following BN3 administration (Figure 3A). Candidate regions were chosen based on their potential roles in anesthesia emergence [15]. Significant c-Fos induction was observed in the medial prefrontal cortex (mPFC), paraventricular thalamus (PVT), and ventral tegmental area (VTA) (Figure 3B, mPFC: *t*_(4)_ = 3.81, *p*<0.05; PVT: *t*_(4)_ = 3.78, *p*<0.05; VTA: *t*_(4)_ = 9.31, *p*<0.001). To determine whether activity in these regions was necessary for BN3’s effects, we performed chemogenetic inhibition (Figure 3C). AAV-hM4Di was injected into the mPFC, PVT, or VTA. Three weeks later, clozapine-N-oxide (CNO; 3 mg/kg, i.p.) was administered 30 min before propofol and BN3. Inhibition of VTA neurons significantly blocked BN3’s emergence-promoting effects (Figure 3D, *t*_(8)_ = 2.67, *p*<0.05). Inhibition of the mPFC also appeared to delay BN3-induced recovery, though this effect did not reach statistical significance (Figure 3D, *t*_(8)_ = 2.00, *p* = 0.08). In contrast, inhibition of the PVT had minimal impact (Figure 3D, *t*_(8)_ = 0.51, *p* = 0.63). As a control, CNO alone did not alter the recovery time from propofol anesthesia by BN3 (Supplementary figure 5).

**Figure 3.**
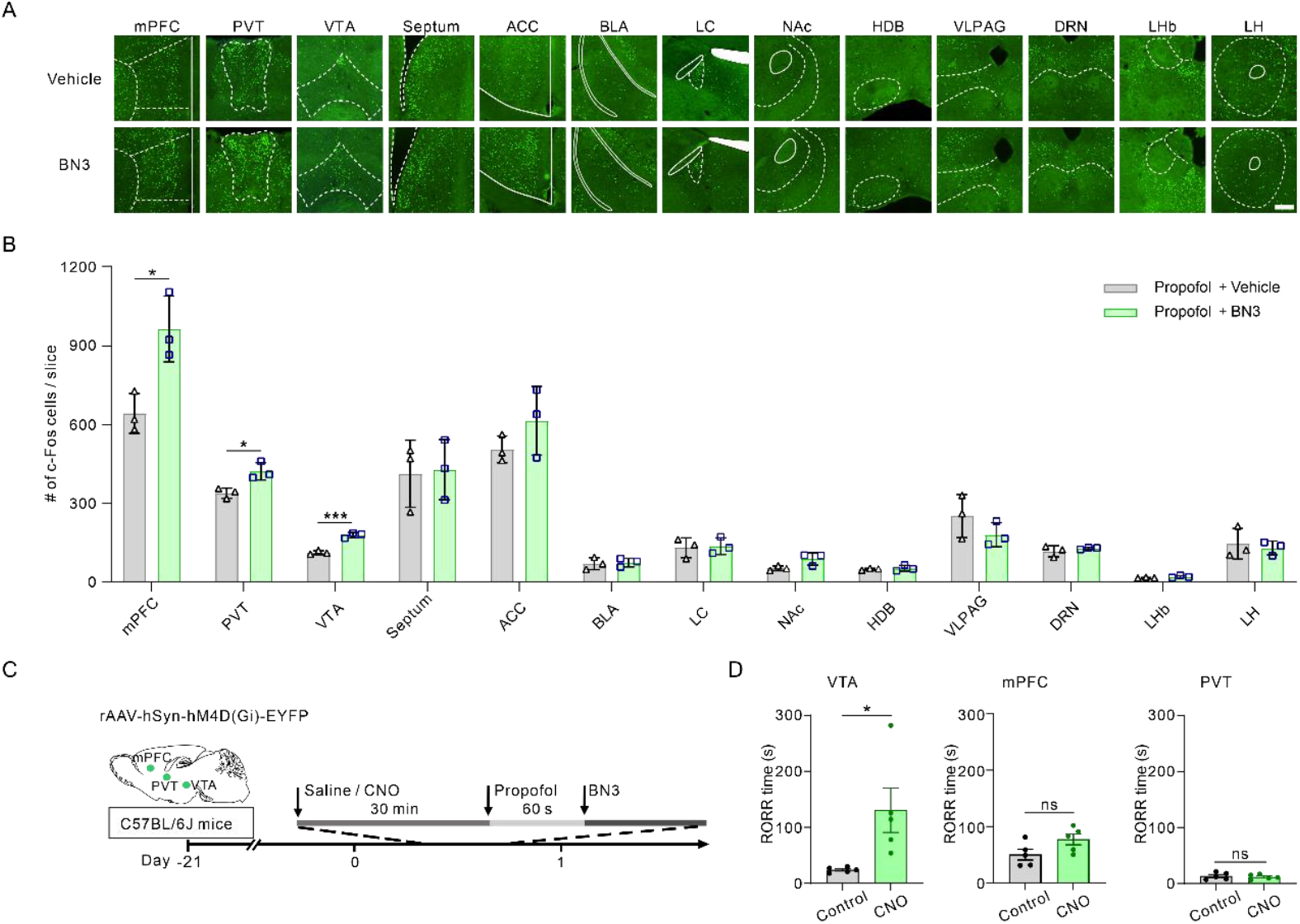
BN3 activates Ventral tegmental area (VTA) to promote emergence from anesthesia. (A) Representative confocal microscopy images showing c-Fos immunoreactivity (green) in coronal brain sections from mice treated with vehicle (top) or BN3 (bottom). Scale bar: 100 µm. mPFC (Medial prefrontal cortex), PVT (Paraventricular thalamus), VTA (Ventral tegmental area), ACC (Anterior cingulate cortex), BLA (Basolateral amygdala), LC (Locus coeruleus), HDB (Horizontal limb of the diagonal band), VLPAG (Ventrolateral periaqueductal gray), DRN (Dorsal raphe nucleus), LHb (Lateral habenula), LH (Lateral hypothalamus). (B) Quantification of c-Fos^+^ cells in the brain regions shown in (A). BN3 significantly increased neuronal activation in the mPFC, PVT and VTA. Data are represented as mean number of c-Fos^+^ cells averaged across three slices from each mouse, n = 3 mice per group. Unpaired two-tailed *t*-test, \**p* < 0.05, \*\*\**p* < 0.001. (C) Schematic of the chemogenetic inhibition protocol. Mice were injected with inhibitory DREADDs (hM4Di) in targeted brain areas 21 days before the experiment. At the day of experiment, CNO was intraperitonially injected 30 minutes prior to propofol anesthesia and BN3 administration. (D) Chemogenetic inhibition of VTA blunted the emergence-accelerating effect of BN3. Return of the righting reflex (RORR) times are shown for control (Saline + BN3) and inhibition (CNO + BN3) groups. n = 5 mice per group. Unpaired two-tailed *t*-test, \**p* < 0.05.

The VTA has recently been identified as a key structure for regulating sleep and wakefulness [16] and has also been implicated in playing a critical role in the emergence from anesthesia [17, 18]. To further test the involvement of VTA in promoting emergence from anesthesia by BN3, AAV-GCaMP6s was injected into the VTA of mice (Figure 4A). Following propofol administration, intravenous BN3 rapidly increased VTA neuronal activity, as evidenced by enhanced GCaMP signals immediately after BN3 injection (Figure 4B-D, *t*_(8)_ = 6.09, *p*<0.001), whereas this effect was abolished by CNO in hM4Di-expressing mice (Figure 4E-H, *t*_(8)_ = 3.00, *p*< 0.05). EEG recordings showed that BN3 decreased delta activity while increasing beta and gamma activity (Figure 4I-L, Delta: *t*_(2)_ = 7.93, *p*<0.05; Beta: *t*_(2)_ = 5.67, *p*<0.05; Gamma: *t*_(2)_ = 4.35, *p*<0.05). These EEG changes were largely blocked by chemogenetic inhibition of VTA (Figure 4M-P).

Taking together, these results identify VTA as a key brain region mediating BN3’s emergence-promoting effects.

**Figure 4.**
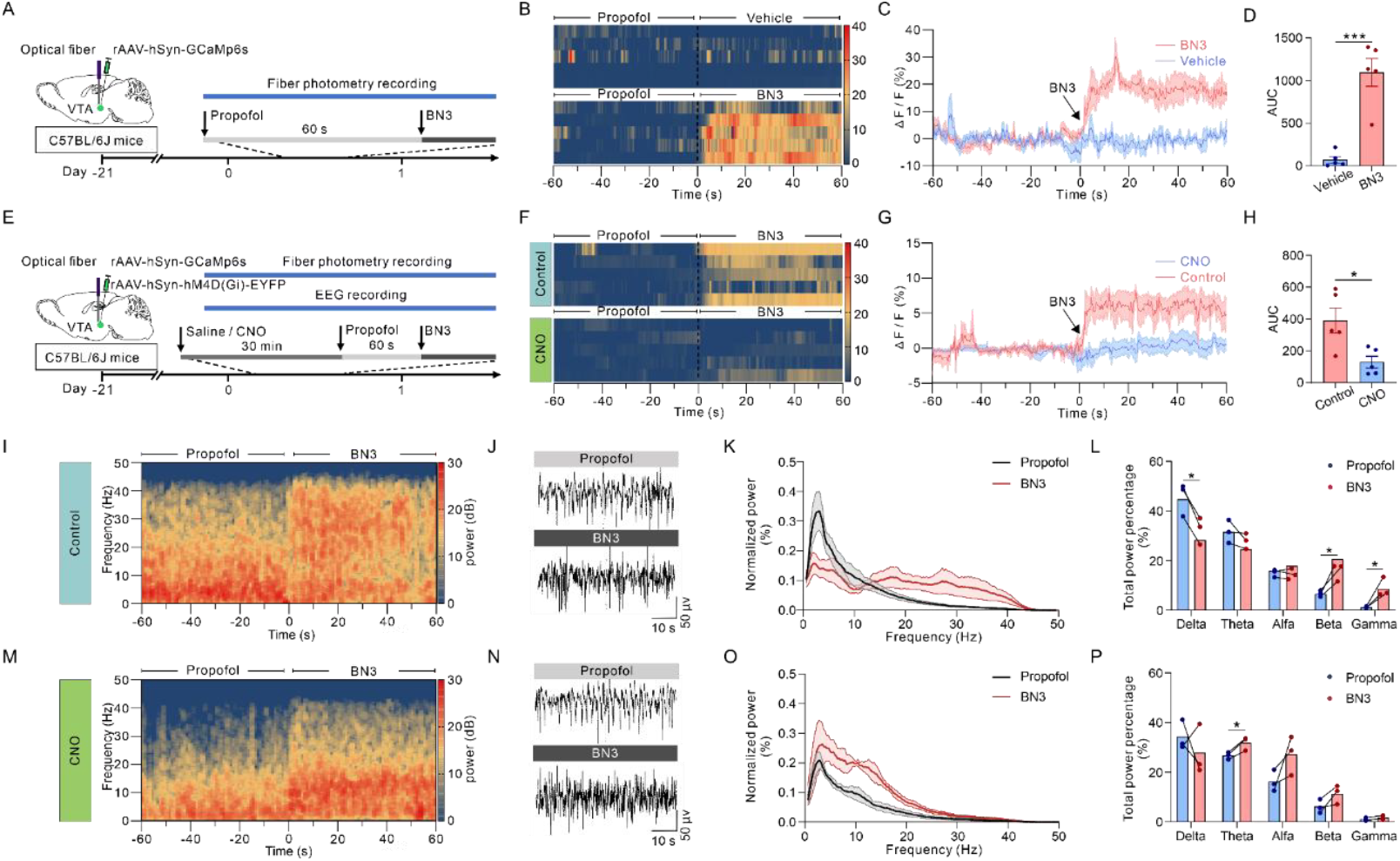
VTA neuronal activity is critical for BN3’s emergence-promoting effect. (A) Schematic illustrating of the fiber photometry protocol. (B) Population heatmap of normalized GCaMp6s fluorescence signals in the VTA, aligned to BN3 administration (time = 0). Each row represents one mouse. BN3 evoked a rapid and sustained increase in calcium activity. (C) Representative averaged ΔF/F0 traces of VTA GCaMp6s signals from individual mice in the vehicle (blue) and BN3 (red) groups. (D) Quantification the area under the curve (AUC) for the 60 seconds period following BN3/vehicle administration; n = 5 mice per group. Unpaired two-tailed *t*-test, \*\*\**p* < 0.001. (E–H) Chemogenetic inhibition of VTA neurons attenuated the BN3-induced calcium response. (E) Schematic illustrating experimental protocol. (F) Population heatmap, (G) representative traces, and (H) AUC quantification for mice expressing hM4Di in VTA neurons; n = 5 mice per group. Unpaired two-tailed *t*-test, \**p* < 0.05. (I–L) BN3 altered cortical EEG signatures toward an awake-like state. (I) Time-frequency spectrogram (heatmap) of cortical EEG power before and after BN3 administration. (J) Representative raw EEG traces under deep propofol anesthesia (top) and after BN3 administration (bottom). (K) Normalized power plots for the pre- and post-BN3 periods. (L) Quantification of the relative power in Delta (0.5–4 Hz), Theta (4–8 Hz), Alfa (8-15 Hz), Beta (15-25 Hz), and Gamma (25-50 Hz) frequency bands. BN3 significantly decreased delta power and increased beta and gamma power; n = 3 mice. Paired two-tailed *t*-test, \**p* < 0.05. (M–P) Chemogenetic inhibition of VTA neurons blunted BN3-induced EEG changes. (M– P) Parallel analyses to panels I-L in mice with VTA inhibition (CNO + BN3 group). The spectral shift toward wakefulness was abolished; n = 3 mice. Paired two-tailed *t*-test, \**p* < 0.05.

### 4. BN3 Accelerates Anesthesia Recovery by Activating Glutamate Neurons in the VTA

The VTA contains three major neuronal subtypes: dopamine, GABA, and glutamate neurons [19]. We next sought to determine which neuronal subtype mediates BN3’s effects. Brains were obtained after BN3 administration (Figure 5A). Co-staining for c-Fos and neuronal markers using fluorescence in situ hybridization revealed the significant increase of co-localization between c-Fos and glutamatergic marker following BN3 administration (Figure 5B-C, *t*_(10)_ = 3.24, *p*<0.01). No significant change of overlap was observed with either GABAergic or dopaminergic markers.

**Figure 5.**
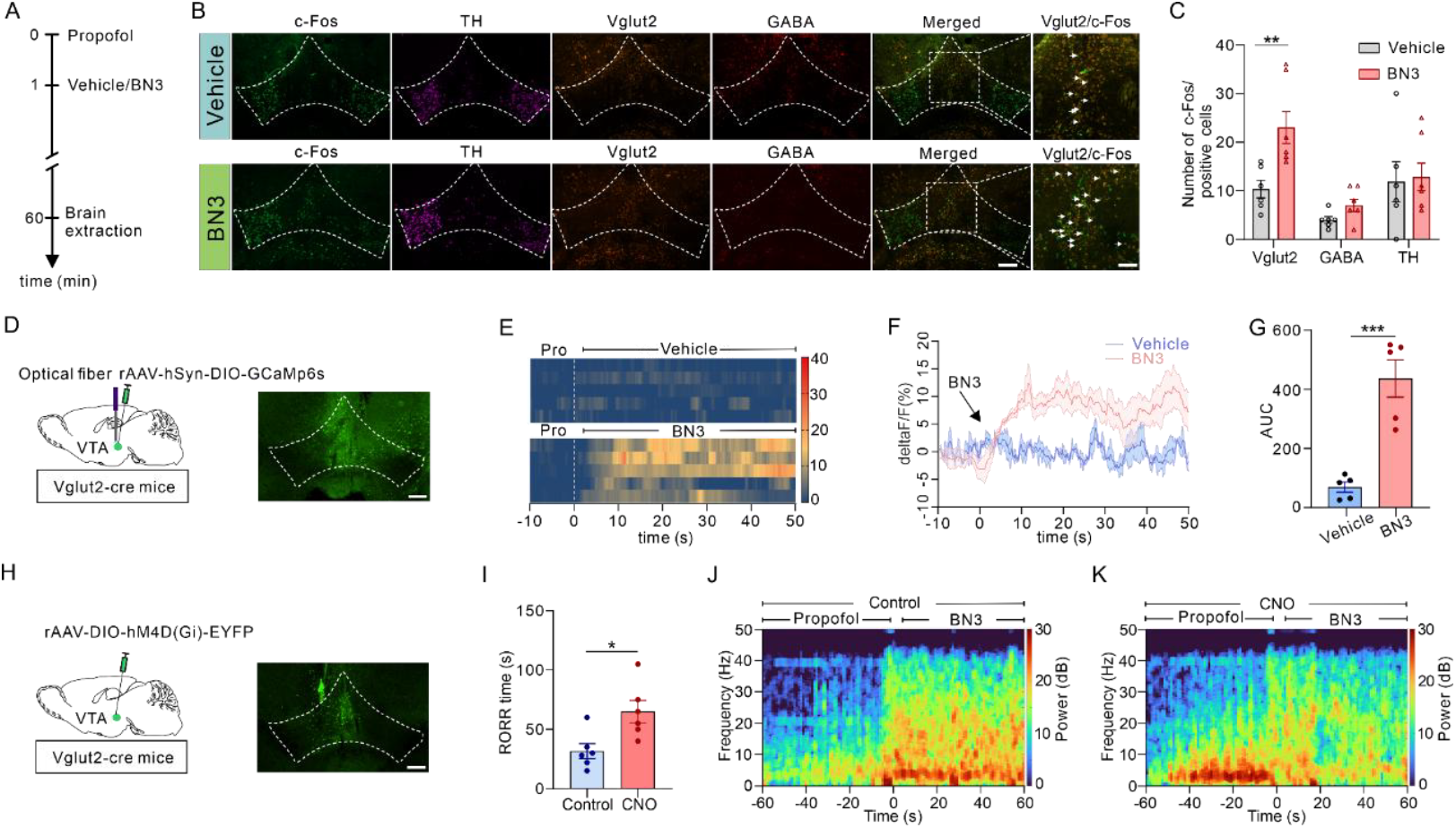
Glutamatergic neurons in the VTA mediate the emergence-promoting effect of BN3. (A) Schematic of the experimental timeline. (B) Representative fluorescence in situ hybridization (FISH) images of coronal VTA sections from BN3-treated mice, showing *c-Fos* mRNA (green) with mRNA markers for dopaminergic (*TH*, purple), glutamatergic (*Vglut2*, yellow), GABAergic (*GABA*, red) neurons. Merged images showing co-localization of *Vglut2* with *c-Fos*. Scale bar: 200 µm, 50 µm in zoom in images. (C) Quantification of activated neuronal subtypes. The number of *Vglut2*^*+*^ neurons that were c-fos^+^ was significantly higher in BN3 group than vehicle group; n = 6 mice. Unpaired two-tailed *t*-test, \*\**p* < 0.01. (D) Schematic of AAV virus injection and optic fiber implantation in the VTA of *Vglut2*-Cre mice to express GCaMP6s selectively in VTA glutamatergic neurons. Right: Representative histology confirming targeted virus expression. (E) Population heatmap of normalized GCaMP6s signals from VTA glutamatergic neurons, aligned to BN3 administration. (F) Representative calcium traces from individual mice. (G) Quantification of the neural response as the area under the curve (AUC); n = 5 mice per group. Unpaired two-tailed *t*-test, \*\*\**p* < 0.001. (H, I) Chemogenetic inhibition of VTA glutamatergic neurons partially reversed BN3’s effect. (H) Schematic of AAV-mediated expression of the inhibitory DREADD hM4Di in VTA glutamatergic neurons of *Vglut2*-Cre mice. Right: Representative image confirming viral expression. (I) Inhibition (via CNO) of VTA glutamatergic neurons significantly delayed RORR time caused by BN3; n = 6 mice per group. Unpaired two-tailed *t*-test, \**p* < 0.05. (J–K) Chemogenetic inhibition of VTA glutamatergic neurons blunted BN3-induced EEG changes.

To directly assess glutamatergic activity, we expressed GCaMp6s selectively in VTA glutamate neurons using *Vglut2*-Cre mice (Figure 5D). Following propofol administration, intravenous BN3 rapidly increased VTA glutamate neuron activity, as indicated by elevated GCaMP signals (Figure 5E–G, *t*_(8)_ = 5.60, *p*<0.001). To test necessity, AAV-DIO-hM4Di was injected into the VTA of *Vglut2*-Cre mice (Figure 5H). CNO injection significantly prolonged recovery time after BN3 administration (Figure 5I, *t*_(10)_ = 2.94, *p*<0.05) and abolished the BN3-induced EEG changes (Figure 5J-K). Conversely, optogenetic activation of VTA glutamate neurons alone was sufficient to accelerate emergence from propofol anesthesia (Supplementary figure 6).

**Figure 6.**
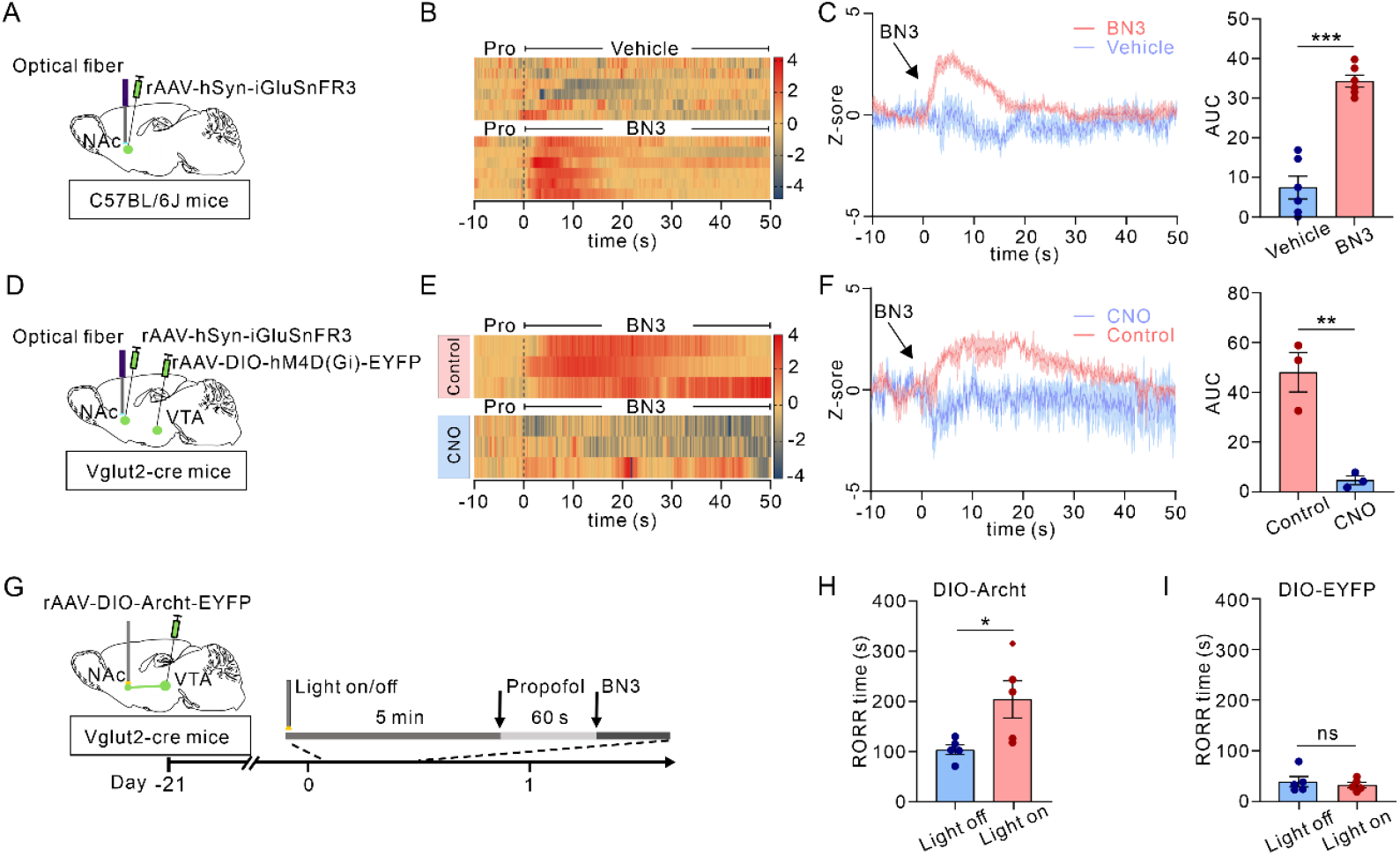
BN3 promotes emergence via VTA glutamatergic projections to the nucleus accumbens (NAc). (A) Schematic for in vivo glutamate sensing in the NAc. (B) Population heatmap of normalized iGluSnFR3 fluorescence signals in the NAc, aligned to BN3 administration (time = 0). Each row represents one animal. (C) Left: Representative traces of glutamate transients in the NAc for vehicle (blue) and BN3 (red) groups. Right: Quantification the area under the curve (AUC) for the 30 seconds following BN3/vehicle administration. BN3 significantly increased glutamate release in the NAc; n = 6 mice per group. Unpaired two-tailed *t*-test, \*\*\**p* < 0.001. (D–F) Chemogenetic inhibition of VTA glutamatergic neurons prevented BN3-evoked glutamate release. (D) Schematic for expressing the inhibitory hM4Di in VTA glutamatergic neurons and glutamate sensor iGluSnFR3 in the NAc. (E) Population heatmap and (F) representative traces with AUC quantification for mice treated with CNO (inhibition) or vehicle prior to 30 seconds following BN3 administration. Inhibition abolished the BN3-induced glutamate transient; n = 3 mice per group. Unpaired two-tailed *t*-test, \*\**p* < 0.01. (G–I) Optogenetic inhibition of VTA→NAc projections blunted BN3’s behavioral effect. (G) Experimental timeline. Archaerhodopsin (Archt) or control (EYFP) was expressed in VTA glutamatergic neurons of *Vglut2*-Cre mice, and an optic fiber was implanted in the NAc. Continuous yellow light (λ = 594 nm, 5-8mW) was delivered 5-min before propofol administration until emergence. (H) Optogenetic inhibition of the VTA→NAc pathway reversed the emergence-accelerating effect of BN3 in Archt-expressing mice. This was not observed in the control animals (I); n = 5 mice per group. Unpaired two-tailed *t*-test, \**p* < 0.05; ns, not significant.

These results demonstrate that activation of VTA glutamate neurons is both necessary and sufficient for BN3’s emergence-promoting effects.

### 5. VTA Glutamate Neurons (VTA^Glu^) promote awaking from anesthesia via nucleus accumbens (NAc) and mPFC

We next sought to identify the downstream targets of VTA glutamate neurons that mediate BN3-induced recovery from general anesthesia.

It has been shown that VTA glutamatergic projection to NAc is important for its wakefulness-promoting effect [16]. We were wondering whether this pathway also participates in emergence-promoting effect of BN3. Because BN3 activates VTA glutamate neurons, we hypothesized that this should enhance glutamate release in downstream targets. Indeed, fiber photometry using the genetically encoded glutamate sensor iGluSnFR3 revealed a significant increase in glutamate release in the NAc following BN3 administration (Figure 6A-C, *t*_(10)_ = 8.30, *p*<0.001). This effect was abolished when VTA glutamate neurons were chemogenetically inhibited (Figure 6D-F, *t*_(4)_ = 5.35, *p*<0.01), confirming VTA glutamate neurons as the upstream source of the BN3-evoked glutamate signal in the NAc. To directly test the involvement of VTA^Glu^ to NAc projections in BN3’ emergence-promoting effect, an inhibitory opsin was specifically expressed in VTA glutamate neurons and optical fiber was placed in the NAc (Figure 6G). When VTA^Glu^ projections to NAc were specifically inhibited, BN3-induced recovery from propofol anesthesia was significantly prolonged (Figure 6H-I, *t*_(8)_ = 2.61, *p*<0.05), indicating that VTAGlu–NAc pathway is required for BN3’s emergence-promoting effects.

VTA glutamate neurons also project to mPFC [20]. Chemogenetic inhibition suggested that mPFC may also contribute to BN3’s effects (Figure 3E). To test this directly, similar experiments were performed as described for the NAc. BN3 administration elicited a robust increase in glutamate release in the mPFC (Figure 7A-C, *t*_(10)_ = 8.82, *p*<0.001), which was prevented by chemogenetic inhibition of VTA glutamate neurons (Figure 7D-F, *t*_(4)_ = 6.06, *p*<0.01). Last, inhibition of VTAGlu– mPFC projections significantly delayed BN3-induced recovery from propofol anesthesia (Figure 7G-I, *t*_(10)_ = 4.56, *p*<0.01).

**Figure 7.**
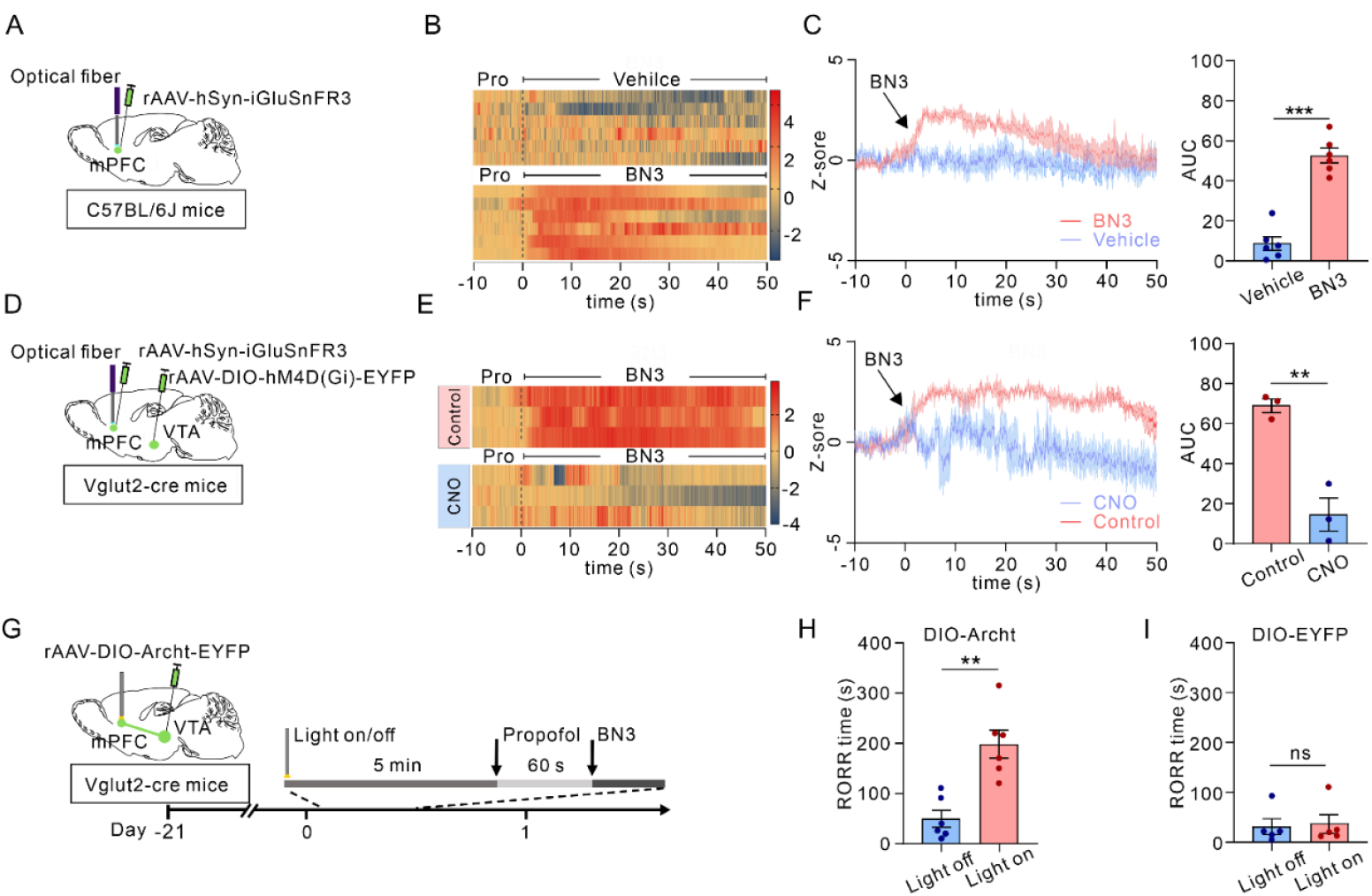
BN3 also promotes emergence via a parallel VTA→medial prefrontal cortex (mPFC) glutamatergic pathway. (A) Schematic for in vivo glutamate sensing in the mPFC. (B) Population heatmap of normalized iGluSnFR3 fluorescence signals in the mPFC, aligned to BN3 administration (time = 0). Each row represents one animal. (C)Left: Representative traces of glutamate transients in the mPFC for vehicle (blue) and BN3 (red) groups. Right: Quantification of the area under the curve (AUC) for the 30 seconds period following BN3/vehicle administration. BN3 potently increased glutamate release in the mPFC; n = 6 mice per group. Unpaired two-tailed *t*-test, \*\*\**p* < 0.001. (D–F) Chemogenetic inhibition of VTA glutamatergic neurons attenuated BN3-evoked glutamate release. (D) Schematic for expressing the inhibitory hM4Di in VTA glutamatergic neurons and glutamate sensor iGluSnFR3 in the mPFC. (E) Population heatmap and (F) Representative traces with AUC quantification for mice pre-treated with CNO (inhibition) or vehicle (control). Inhibition significantly reduced the glutamate transient; n = 3 mice per group. Unpaired two-tailed *t*-test, \*\**p* < 0.01. (G–I) Optogenetic inhibition of VTA→mPFC projections partially reversed BN3’s behavioral effect. (G) Experimental timeline. Archaerhodopsin (Archt) or control (EYFP) was expressed in VTA glutamatergic neurons of *Vglut2*-Cre mice, with an optic fiber in the mPFC. Continuous yellow light (λ = 594 nm, 5-8mW) was delivered 5-min before propofol administration until emergence. (H) Optogenetic inhibition of the VTA→mPFC pathway significantly attenuated the emergence-accelerating effect of BN3 in Archt-expressing mice, but not in control group (I); n = 6 mice per group. Unpaired two-tailed *t*-test, \*\**p* < 0.01; ns, not significant.

Together, these data support that VTA glutamate neurons and their downstream targets underlie the emergence-promoting effects of BN3.

## Discussion

In this study, we identify BN3 (L-(−)-camphor) as a natural-product-derived small molecule that accelerates emergence from general anesthesia and define a circuit mechanism that accounts for its broad efficacy. BN3 shortened recovery from propofol across five mammalian species, including dogs and rhesus monkeys, and remained effective after clinically relevant continuous-infusion paradigms. In rodents, BN3 also facilitated emergence from mechanistically diverse anesthetics, extending beyond intravenous GABAergic hypnotics to include a benzodiazepine-site modulator (remimazolam), sodium γ-hydroxybutyrate, an NMDA receptor antagonist (ketamine), and a volatile anesthetic (isoflurane). By combining EEG state readouts with cell-type- and projection-specific causal perturbations, we show that BN3 promotes a rapid cortical state transition toward wake-like activity and recruits a VTA-centered glutamatergic arousal circuit engaging outputs to the nucleus accumbens (NAc) and medial prefrontal cortex (mPFC). Together, these findings support a circuit-based, anesthetic-independent strategy for facilitating emergence and provide a chemically tractable entry point for translational development.

A central clinical constraint in anesthesia practice is that pharmacologic reversal of unconsciousness remains largely limited to drug-class–specific antagonists, leaving most recoveries to the decline of anesthetic effect-site concentrations [7]. At the systems level, however, induction and emergence are increasingly viewed as asymmetric state transitions shaped by endogenous arousal networks rather than as simple pharmacokinetic mirror images [8, 15, 21]. Our findings extend this framework by demonstrating a small-molecule intervention whose efficacy generalizes across anesthetics with divergent molecular actions. Several aspects of the data argue against a purely pharmacokinetic explanation in which BN3 accelerates recovery solely by enhancing anesthetic clearance: BN3 retains efficacy after continuous infusion (where anesthetic burden is high), evokes rapid VTA activation tightly time-locked to its administration, and produces a prompt EEG reconfiguration consistent with an active brain-state transition. Moreover, optogenetic activation of VTA glutamatergic neurons recapitulates key features of BN3-induced emergence, supporting a circuit mechanism capable of driving the transition. Definitive pharmacokinetic– pharmacodynamic studies and molecular target identification will nonetheless be essential to establish how BN3 engages this circuitry and to quantify any contribution of altered anesthetic disposition.

Our mechanistic experiments nominate the ventral tegmental area (VTA) as a critical node mediating BN3’s emergence-promoting effect. The VTA has previously been implicated in arousal and reanimation through electrical stimulation [17, 18] and dopaminergic mechanisms [22-24]. Our results converge on this hub while refining the identity of the key effectors. BN3 preferentially activates VTA glutamatergic neurons, as supported by subtype-resolved activity mapping and fiber photometry, and chemogenetic inhibition of this population markedly blunts both the behavioral acceleration of emergence and the associated wake-like EEG spectral shift. Conversely, direct optogenetic activation of VTA glutamatergic neurons is sufficient to promote emergence under propofol anesthesia. These observations indicate that fast excitatory signaling within midbrain arousal circuitry can serve as a potent lever for state transition, and they position VTA glutamatergic neurons as a mechanistically grounded substrate for broad-spectrum emergence facilitation rather than as a correlate of recovery.

Projection-specific experiments further define a dual-output circuit architecture underlying BN3-induced emergence. BN3 increases glutamate release in two major downstream targets—the NAc and mPFC—and disrupting either VTA→NAc or VTA→mPFC glutamatergic output blunts BN3’s behavioral effect. These two targets are well positioned to contribute complementary components of awakening: the NAc integrates motivational salience with state regulation and can gate behavioral arousal [25, 26], whereas the mPFC is a key node in networks supporting cortical activation and conscious-level control [27]. One parsimonious model is therefore that BN3 recruits a coordinated VTA glutamatergic program in which striatal output pathways help trigger the transition while prefrontal circuitry helps stabilize a wake-like cortical regime. Importantly, VTA glutamatergic neurons are heterogeneous, and additional downstream nodes may shape emergence depending on anesthetic class, depth of anesthesia, and physiological context. Dissecting projection-defined subpopulations, and determining how these outputs interact with other canonical arousal systems (including basal forebrain, thalamocortical loops, and neuromodulatory pathways), will be important for refining a circuit-level account of anesthetic emergence.

From a translational perspective, an emergence-promoting strategy that does not require matching each anesthetic’s molecular target could address clinically important problems such as delayed emergence and heterogeneous recovery trajectories, particularly in infusion-based practice patterns. The cross-species efficacy of BN3 for propofol, including in non-human primates, provides an initial bridge toward clinical feasibility, while the comparison with flumazenil highlights the conceptual advantage of a circuit-recruiting agent over class-restricted antagonism. At the same time, translation will require rigorous definition of the pharmacokinetic–pharmacodynamic relationship, development of a scalable intravenous formulation, and systematic assessment of cardiorespiratory and neurobehavioral safety across dosing windows. Because VTA circuits participate in motivation, salience processing, and affective behaviors, future work should explicitly examine whether circuit recruitment can be tuned to promote timely emergence without provoking agitation, dysphoria, or maladaptive arousal states. Finally, while loss and return of the righting reflex together with EEG spectral transitions provide operationally tractable readouts of emergence, the clinical value of an emergence agent will ultimately depend on the quality of recovery—cognitive trajectory, delirium risk, and functional outcomes—particularly in vulnerable populations and in multimodal anesthetic regimens.

In summary, we identify BN3 as a natural-product-derived emergence agent and establish a VTA-centered glutamatergic circuit mechanism that can be pharmacologically recruited to facilitate anesthetic-independent emergence. By linking broad efficacy to a causally specified arousal circuit and its downstream effectors, this work reframes emergence as an actively controllable process and provides a foundation for circuit-targeted strategies to improve recovery and perioperative safety.

## Acknowledgments

This work was supported by the National Natural Science Foundation of China (82425054 and 82273784 to B.K.), Brain Science and Brain-like Intelligence Technology-National Science and Technology Major Project (2025ZD0214904 to B.K.), Science and Technology Department of Sichuan Province (2024NSFSC0048 to B.K.), 1.3.5 Project for Disciplines of Excellence, West China Hospital, Sichuan University (ZYGD25002 and ZYGD23025 to B.K.).

