## Supplementary materials for "Pharmacological recruitment of a VTA glutamatergic arousal circuit by the natural product BN3 drives broad-spectrum emergence from general anesthesia"

### Supplementary table 1. Natural product compound library

| Compd. No | CAS | NAME | SOURCE |
|---|---|---|---|
| BN1 | 87-44-5 | β-Caryophyllene | Bidepharm |
| BN2 | 89-48-5 | (-)-Menthyl Acetate- | Bidepharm |
| BN3 | 464-48-2 | (-)-Camphor | Bidepharm |
| BN4 | 117-39-5 | Quercetin | Bidepharm |
| BN5 | 482-36-0 | Hyperoside | Bidepharm |
| BN6 | 520-26-3 | Hesperidin | Bidepharm |
| BN7 | 60-33-3 | Linoleic acid | Bidepharm |
| BN8 | 110-15-6 | Succinic acid | Bidepharm |
| BN9 | 57-10-3 | Palmitic acid | Bidepharm |
| BN10 | 104-54-1 | Cinnamyl alcohol | Bidepharm |
| BN11 | 100-51-6 | Benzyl alcohol | Bidepharm |
| BN12 | 60-12-8 | Phenethyl alcohol | Bidepharm |
| BN13 | 520-34-3 | Diosmetin | Bidepharm |
| BN14 | 5957-80-2 | Carnosol | Bidepharm |
| BN15 | 520-36-5 | Apigenin | Bidepharm |
| BN16 | 142-26-7 | 2-Acetylaminoethanol | Bidepharm |
| BN17 | 970-74-1 | (-)-Epigallocatechin | Bidepharm |
| BN18 | 77-52-1 | Ursolic acid | Bidepharm |

| | | | |
|---|---|---|---|
| BN19 | 1257-08-5 | (-)-Epicatechin gallate | Bidepharm |
| BN20 | 2883-98-9 | alpha-Asarone | Bidepharm |
| BN21 | 470-67-7 | 1,4-Cineole | Bidepharm |
| BN22 | 89-78-1 | DL-Menthol | Bidepharm |
| BN23 | 81-25-4 | Cholic acid | Bidepharm |
| BN24 | 10458-14-7 | Menthone | Bidepharm |
| BN25 | 58-55-9 | Theophylline | Bidepharm |
| BN26 | 83-67-0 | Theobromine | Bidepharm |
| BN27 | 65-85-0 | Benzoic acid | Bidepharm |
| BN28 | 327-97-9 | Chlorogenic acid | Bidepharm |
| BN29 | 50-28-2 | β-Estradiol | Bidepharm |
| BN30 | 490-46-0 | L-Epicatechin | Bidepharm |
| BN31 | 76-49-3 | Bornyl acetate | Bidepharm |
| BN32 | 57-88-5 | Cholesterol | Bidepharm |
| BN33 | 327-97-9 | Chlorogenic acid | Bidepharm |
| BN34 | 110-17-8 | Fumaric acid | Bidepharm |
| BN35 | 1135-24-6 | Ferulic Acid | Bidepharm |
| BN36 | 93-15-2 | Methyl eugenol | Bidepharm |
| BN37 | 485-72-3 | Formononetin | Bidepharm |
| BN38 | 5273-86-9 | β-Asarone | Bidepharm |
| BN39 | 20575-57-9 | Calycosin | Bidepharm |

| | | | |
|---|---|---|---|
| BN40 | 63569-07-3 | Microhelenin C | Bidepharm |
| BN41 | 16503-32-5 | Brevilin A | Bidepharm |
| BN42 | 491-70-3 | Luteolin | Bidepharm |
| BN43 | 487-11-6 | Elemicin | Bidepharm |
| BN44 | 607-80-7 | Sesamin | Bidepharm |
| BN45 | 133-05-1 | (-)-ASARININ 97 | Bidepharm |
| BN46 | 535-83-1 | Trigonelline | Bidepharm |
| BN47 | 121-34-6 | Vanillic acid | Bidepharm |
| BN48 | 99-50-3 | Protocatechuic acid | Bidepharm |
| BN49 | 123-35-3 | Myrcene | Bidepharm |
| BN50 | 78-70-6 | Linalool | Bidepharm |
| BN51 | 1139-30-6 | Caryophyllene oxide | Bidepharm |
| BN52 | 487-36-5 | (+)-PinoresinolL | Bidepharm |
| BN53 | 98-55-5 | α-Terpineol | Bidepharm |
| BN54 | 483-76-1 | (+)-δ-cadinene | Bidepharm |
| BN55 | 494-90-6 | Menthofuran | Bidepharm |
| BN56 | 562-74-3 | Terpinen-4-ol | Bidepharm |
| BN57 | 515-13-9 | (-)-β-Elemene | Bidepharm |

All compounds were formulated at a working concentration of 50 mM and delivered intravenously as a fixed volume of 0.2 mL per mouse.

### Supplementary figures

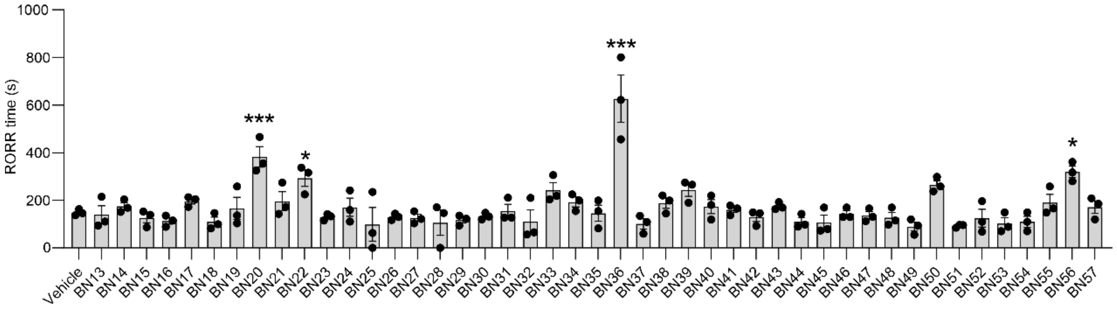


**Supplementary figure 1. Phenotypic screening natural compounds BN13–BN57.**

Screening of natural compounds (BN13–BN57) for their ability to affect the time to emergence after a standardized bolus dose of propofol in mice. BN20, BN22, BN36 and BN56 significantly prolonged the righting reflex recovery (RORR) time induced by propofol, n = 3 mice per group. One-way ANOVA, $*p < 0.05$, $***p < 0.001$.

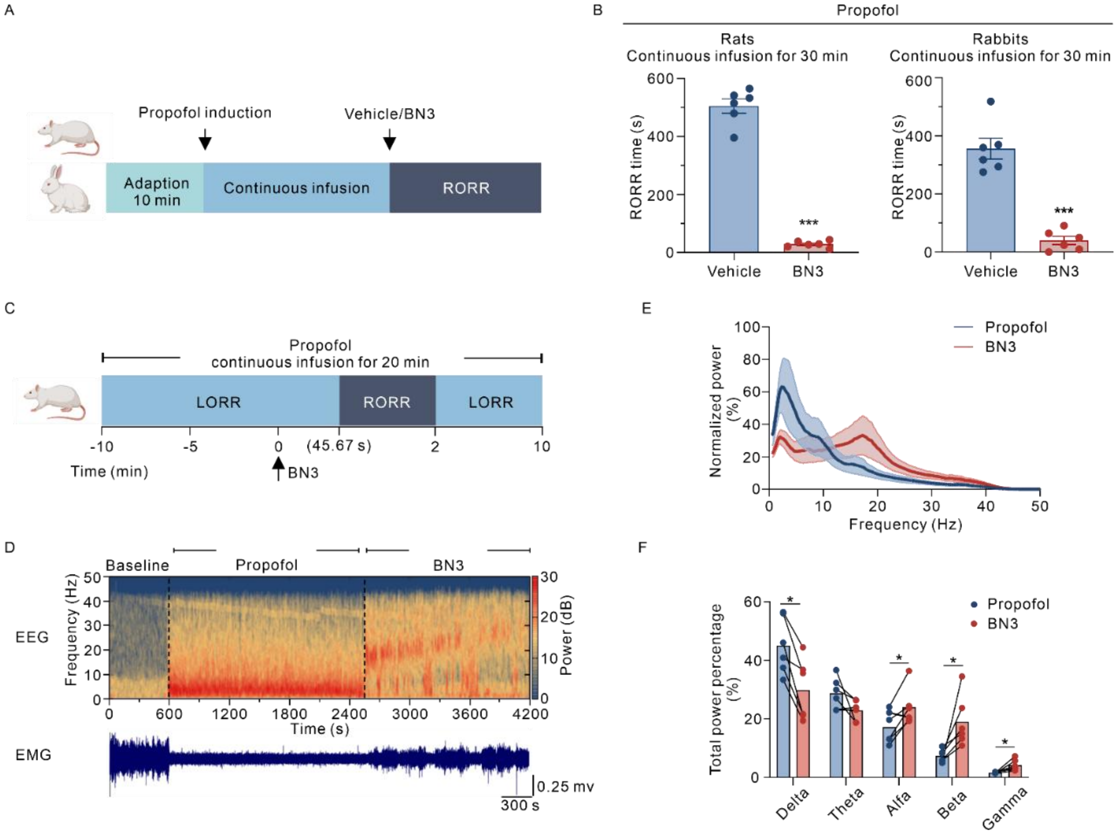


**Supplementary figure 2. BN3 accelerates emergence from continuous propofol anesthesia and restores wake-like brain activity.**

(A) Schematic timeline of the continuous intravenous propofol infusion protocol. Mice received a 30-min infusion of propofol. Vehicle or BN3 was administered at the cessation of infusion, and the time to return of the righting reflex (RORR) was recorded.

(B) BN3 significantly reduced the time to RORR following continuous propofol anesthesia. n = 6 animals per group. Unpaired two-tailed *t*-test, ***p < 0.001.

(C) BN3 induced return of the righting reflex during continuous propofol anesthesia. n = 3 rats, (averaged RORR = 45.67 s).

(D) Time-frequency spectrogram (heatmap) of cortical electroencephalogram (EEG) power before and after BN3 administration, aligned to the time of injection (dashed line). Corresponding EMG signals was shown at the bottom.

(E) Normalized power plots for the 30-minute epochs immediately before (blue) and after (red) BN3 administration.

(F) Quantification of relative power in Delta (0.5–4 Hz), Theta (4–8 Hz), Alfa (8-15 Hz), Beta (15-25 Hz), and Gamma (25-50 Hz) frequency bands. BN3 significantly decreased delta power and increased alfa, beta, and gamma power. Individual paired data points for each rat are connected by lines; n = 6 rats. Paired two-tailed *t*-test, *$p < 0.05$.

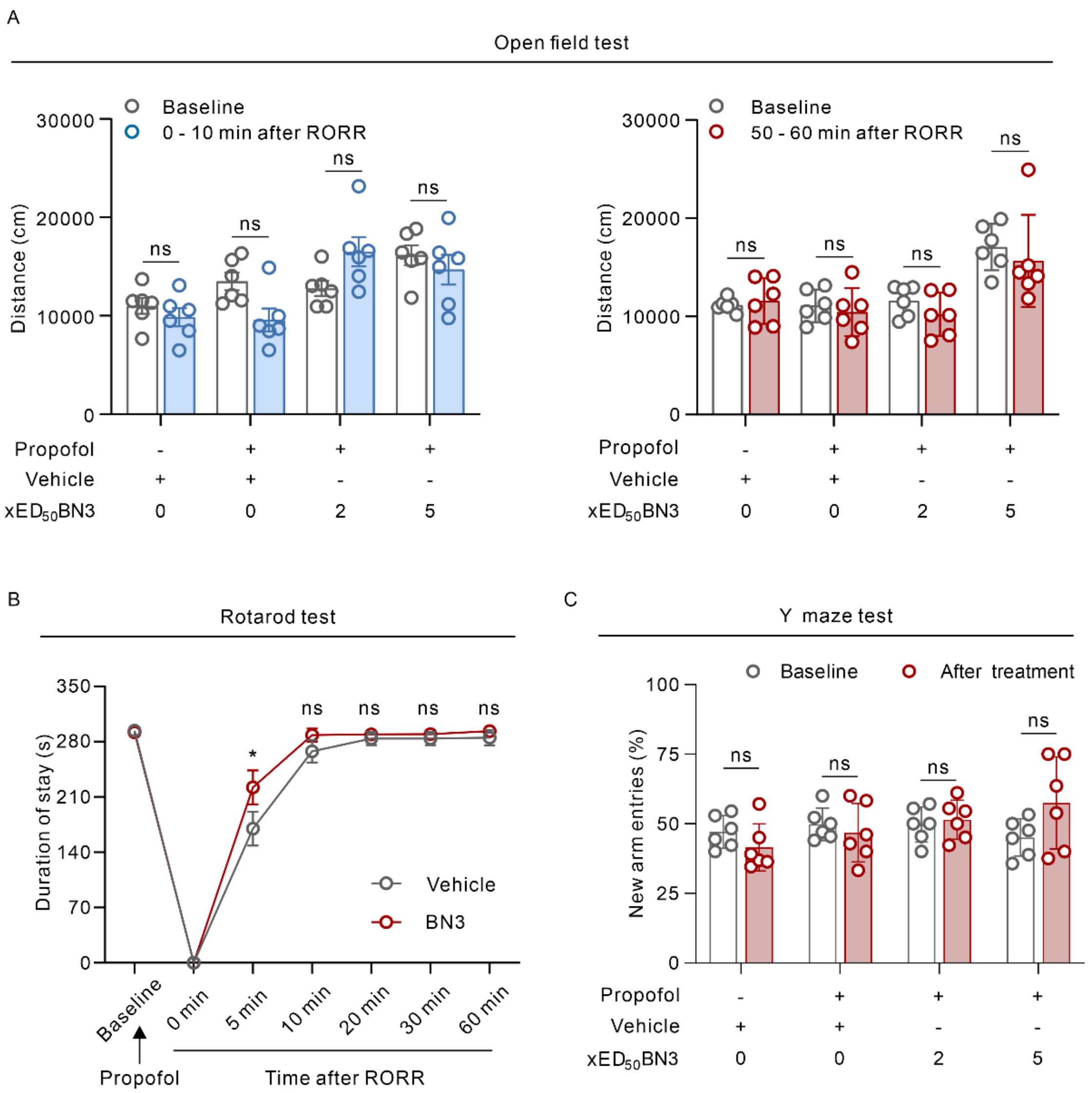


**Supplementary figure 3. Behavioral assessment of mice after emergence from propofol anesthesia indicates a favorable safety profile for BN3.**

(A) Locomotor activity: Mice were allowed to recover from propofol anesthesia and subjected to open field test immediate and 50 minutes after the return of the righting reflex (RORR). Total distance traveled in a 10-minute open field test showed no significant difference between baseline and treatment; n = 6 mice per group. Unpaired two-tailed *t*-test, ns, not significant.

(B) Motor coordination and strength: Latency to fall on an accelerating rotarod. BN3-treated mice performed better or comparably to vehicle controls, indicating no impairment in motor function; n = 6 mice per group. Unpaired two-tailed *t*-test, *$p < 0.05$; ns, not significant.

(C) Spatial recognition memory: Mice were allowed to recover from propofol anesthesia and subjected to Y-maze test when mice resumed normal activity. Percentage of new arm entries were presented. No significant difference was observed between baseline and treatment, suggesting that BN3 treatment did not affect spatial recognition memory; n = 6 mice per group. Unpaired two-tailed *t*-test, ns, not significant.

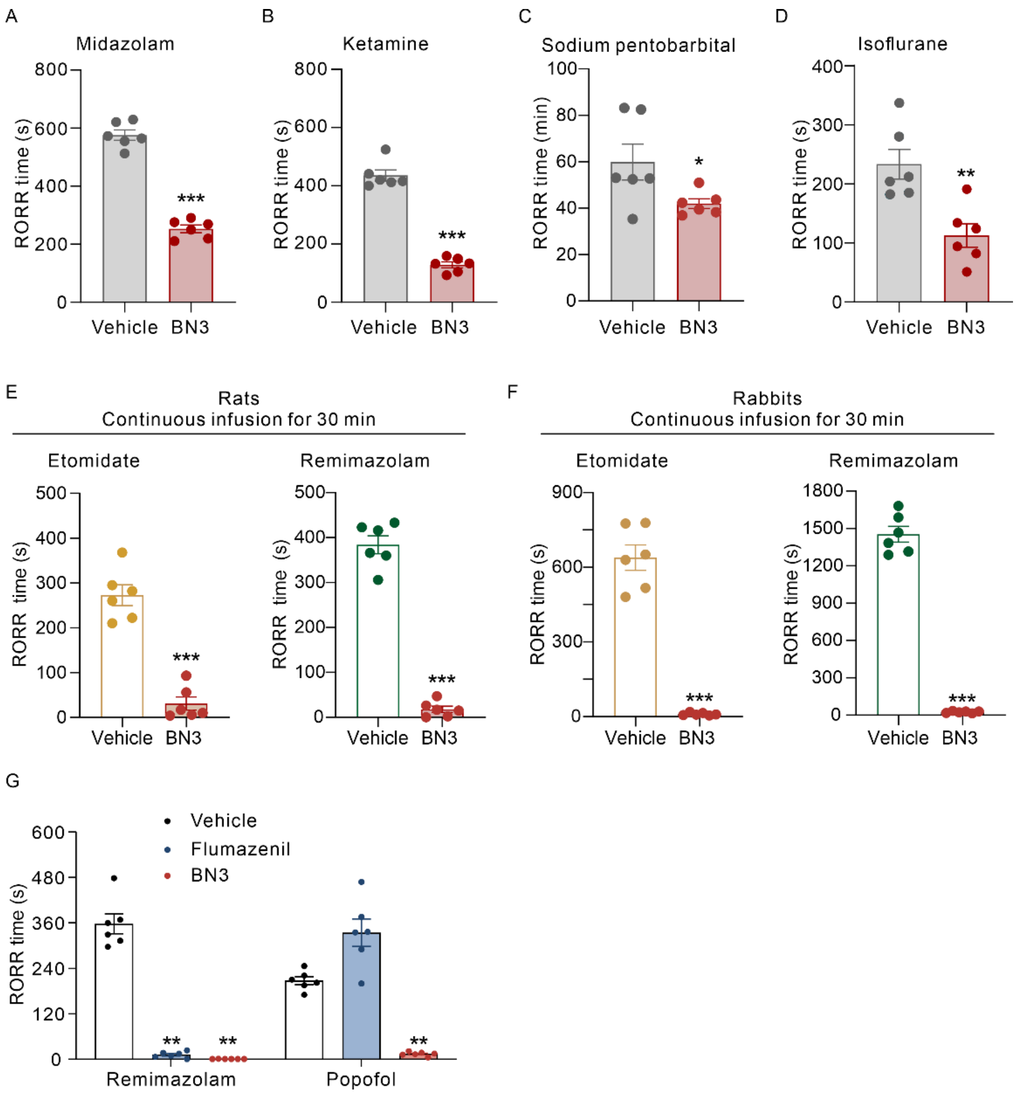


**Supplementary figure 4. BN3 accelerates emergence from diverse anesthetics across species.**

(A) Benzodiazepine agonist: BN3 significantly reduced emergence time from midazolam-induced general anesthesia in mice.

(B) NMDA receptor antagonist: BN3 significantly reduced emergence time from ketamine-induced general anesthesia in mice.

(C) $GABA_A$ receptor: BN3 significantly reduced emergence time from Sodium pentobarbital-induced general anesthesia in mice.

(D) Inhalation anesthetics: BN3 significantly reduced the emergence time in mice following 30 min of general anesthesia with 1.5% isoflurane; n = 6 mice per group. Unpaired two-tailed *t*-test, *$p < 0.05$, **$p < 0.01$, ***$p < 0.001$.

(E, F) Continuous infusion models: BN3 significantly reduced emergence time in rats (E) and rabbits (F) following a continuous intravenous infusion of etomidate or remimazolam; n = 6 animals per group. Unpaired two-tailed *t*-test, ***$p < 0.001$.

(G) Comparison to flumazenil: The emergence-accelerating efficacy of BN3 is different from that of the benzodiazepine antagonist flumazenil. Bar graphs compare RORR times following remimazolam (left) or propofol (right) anesthesia. Flumazenil was only effective in reversing remimazolam, while BN3 significantly accelerated emergence from both anesthetics; n = 6 mice per group. One-way ANOVA, **$p < 0.01$.

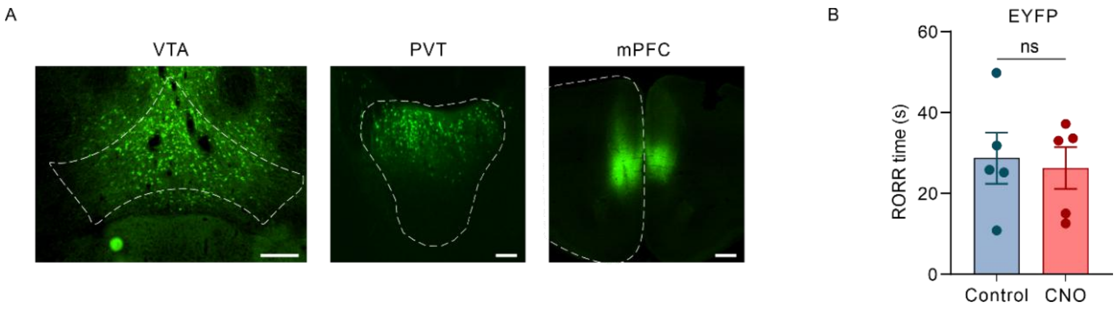


**Supplementary figure 5. Histological verification of viral targeting and a control experiment for chemogenetic inhibition.**

(A) Confocal microscopy images validating targeted viral expression in the VTA (Ventral tegmental area) and PVT (Paraventricular thalamus), Scale bars: 200 µm; mPFC (Medial prefrontal cortex), Scale bar: 500 µm.

(B) Control experiment confirming that the DREADD actuator clozapine N-oxide (CNO) itself does not affect BN3's efficacy. Mice expressing an inert fluorescent protein (control virus) in the VTA were administered CNO (or vehicle) prior to BN3 treatment under standard propofol anesthesia. No significant difference was observed in return of the righting reflex (RORR) times between the CNO and vehicle control groups; n = 5 mice per group. Unpaired two-tailed *t*-test, ns, not significant.

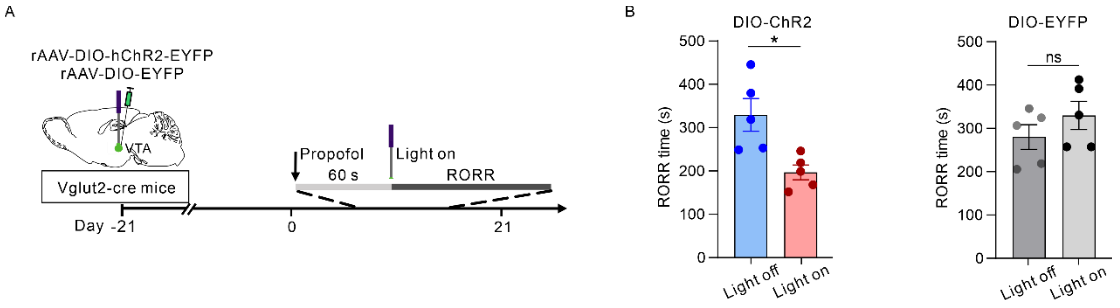


**Supplementary figure 6. Direct optogenetic activation of VTA glutamatergic neurons is sufficient to accelerate emergence from propofol anesthesia.**

(A) Schematic illustrating of experimental protocol. In *Vglut2*-Cre mice, a Cre-dependent channelrhodopsin (AAV-DIO-hChR2-EYFP) or control fluorophore (AAV-DIO-EYFP) was expressed in the VTA. One minute after propofol administration, VTA glutamatergic neurons were activated by pulsed blue light ($\lambda = 473$ nm, 2-5mW, 15-ms pulses, 20Hz) delivered via an implanted optic fiber. The time to return of the righting reflex (RORR) was measured.

(B) Optogenetic activation of VTA glutamatergic neurons significantly reduced RORR time compared to no-light controls in ChR2-expressing mice, but not in EYFP control mice, n = 5 mice per group. Unpaired two-tailed $t$-test, $*p < 0.05$; ns, not significant.
